# Differential adenosine signaling and effects of acute caffeine exposure on alternative stress coping styles in zebrafish (*Danio rerio*)

**DOI:** 10.64898/2026.01.20.700609

**Authors:** Sydney E. Klucas, Ryan Y. Wong

**Affiliations:** Department of Biology, University of Nebraska at Omaha

**Keywords:** Caffeine, Zebrafish, Adenosine pathway, Anxiety, Stress coping styles

## Abstract

Changes within neurotransmitter systems are associated with variation in anxiety-related behavior. The adenosine signaling pathway has been associated with anxiety and caffeine has been utilized as a modulator. However, studies have not considered the impact of an individual’s stress coping style (e.g. proactive, reactive) and corresponding differences in neuromolecular signaling that can influence the behavioral responses. To assess the role of adenosine signaling, we acutely treated reactive and proactive zebrafish with 50 mg/L caffeine and evaluated anxiety-like behavior using a novel tank diving test (NTDT). We then quantified whole-brain gene expression of genes representing distinct parts of the adenosine signaling pathway: adenosine receptors A1B, A2Aa, A2Ab, and A2B (*adora1b, adora2aa, adora2ab*, and *adora2b*, respectively) and enzymes adenosine deaminase (*ada*) and ecto-5’-nucleotidase (*nt5e*). We found significant main effects of coping style, sex, treatment, and coping style by sex by treatment interaction effect on stress behaviors. Specifically, compared to controls, caffeine reduced stress behavior in only reactive males. We also observed significant differential baseline gene expression within the adenosine signaling pathway between the reactive and proactive strains, where reactive zebrafish expressed higher levels of adenosine receptors A1B, A2Ab, A2B, and adenosine deaminase and lower levels of adenosine receptor A2Aa than proactive zebrafish. These findings indicate that variation in adenosine signaling between the stress coping styles and sexes may be contributing to differences in anxiety-related behavior.

## Introduction

Animals encounter many stressful events such as trying to obtain resources amongst competitors or to avoid being prey and often utilize strategies to successfully overcome those stressors. The resulting physiological and behavioral changes to a stressor are mediated in part by the sympathetic-adrenal medullary and hypothalamic-pituitary-adrenal axes^1,2^. Exaggerated or inappropriate responses to stressors are symptoms of anxiety-related disorders, and it is hypothesized that dysregulation of neurotransmitter signaling pathways can facilitate the progression of the disorders^3^. Specifically, variation in the adenosine neurotransmitter system may increase susceptibility to anxiety disorders^4,5^.

Adenosine is a neuromodulator that has a crucial role in cellular signaling. Within the adenosine pathway of the central nervous system, adenosine is an inhibitory neurotransmitter that binds to the A1, A2A, A2B, and A3 adenosine receptors. As G protein-coupled receptors (GPCRs), adenosine A1 and A3 receptors are coupled to G_i/o_ proteins (inhibitory) while adenosine A2A and A2B receptors are coupled to G_s/q_ proteins, causing inhibitory or excitatory effects on several downstream signaling pathways^6,7^. The binding of adenosine to these receptors is essential for the sleep-wake cycle, immune responses, inflammation, and several other downstream effects^8,9^. Adenosine levels are highly regulated through degradation and production by multiple enzymes. Two of the main extracellular enzymes within the pathway include ecto-5’-nucleotidase (nt5e, also known as CD73), which produces adenosine by hydrolyzing adenosine monophosphate (AMP), and adenosine deaminase, which catalyzes the deamination of adenosine into inosine. The adenosine signaling pathway has been implicated in anxiety, wherein both adenosine concentration and adenosine receptor interactions are altered^10–18^. Adenosine levels increase in the brain after chronic stress^10^. Both ecto and cytosolic adenosine deaminase activity decrease following a stressor^10,11,18^, which may contribute to an increase in adenosine levels.

Administration of an adenosine derivative in mice decreased both anxious behavior and adenosine deaminase activity and increased adenosine levels in the cortex and hypothalamus^18,19^. Caffeine is another compound that can alter adenosine signaling through unbiased competitive inhibition of adenosine receptors A1 and A2A. Genetic variation in adenosine receptors A1 and A2A are associated with panic disorder and anxiety following caffeine consumption in humans^12,20–23^, and antagonism of adenosine receptors by caffeine has been found to be implicated in anxiety behavior^13–17,24^. Although different populations of zebrafish and rats differ in their anxiogenic responses^16,17^, it is not yet known if genetic differences in the adenosine pathway are responsible in part for altered anxiety-like behavior.

Zebrafish (*Danio rerio*) are a commonly used animal model for understanding the molecular and pharmacological bases of anxiety due to their ease of maintenance and their highly conserved genetic and physiological features with mammals and other species^25,26^. To facilitate identifying molecular mechanisms of stress- and anxiety-like behaviors, prior studies have utilized zebrafish strains selectively bred to display opposing stress coping styles: proactive and reactive^27–31^. Across different stress assays and time, the reactive zebrafish strain displays higher anxiety-related behaviors and has a faster whole-body cortisol response compared to the proactive zebrafish^27–31^. Further, the two zebrafish strains show distinct baseline neurotranscriptome profiles, including differences in expression within several neurotransmitter pathways, such as dopamine, serotonin, GABA, and adenosine^29,32^. Receptors and enzymes within the zebrafish adenosine pathway are homologous to those of mammals^33^, and their presence in the zebrafish brain is well-documented^34–38^. Within the adenosine pathway, the proactive strain has significantly higher whole-brain expression of adenosine deaminase and ecto-5’-nucleotidase compared to the reactive strain at baseline^29^. In addition to stress coping style, sex differences have been documented in anxiety-related behavior and adenosine signaling, with females generally exhibiting higher anxiety than males across species^24,27,39–42^ and demonstrating sex-specific variation in the adenosine system^43–46^. There is limited understanding of how stress coping styles and sex interact to affect anxiety behavior mediated by adenosine signaling.

In the present study, we investigated the effects of acute caffeine treatment on anxiety-related behavior and gene expression within the adenosine signaling pathway in proactive and reactive strains of zebrafish. We evaluated stress behaviors using the novel tank diving test (NTDT) and quantified whole-brain gene expression of adenosine receptors A1B, A2Aa, A2Ab, and A2B (*adora1b, adora2aa, adora2ab*, and *adora2b*, respectively) as well as the adenosine deaminase (*ada*) and ecto-5’-nucleotidase (*nt5e*) enzymes. We hypothesized that acute caffeine exposure will increase anxiety-like behavior in both coping styles with an amplified anxiogenic response in the reactive strain. At baseline, we predicted that the reactive strain would have increased expression of adenosine receptors and adenosine deaminase compared to the proactive strain. Understanding how an adenosine receptor antagonist impacts adenosine signaling between the two coping styles and sexes will provide insight into a mechanism that may explain differences in their anxiety-related behaviors.

## Methods

### Subjects

We used zebrafish (*Danio rerio*) strains selectively bred to display the proactive or reactive stress coping style. The strains originated from wild-caught zebrafish from Gaighata, India and were then selectively bred based on behavioral response to stress^27^. The two strains have well-documented differences in stress-related responses across several behavioral assays, time, morphology, neuropharmacology, neurotranscriptome analyses, learning and memory, and glucocorticoid responses^27,28,30–32,47–50^. In this study, all fish were 9-11 months old when testing began and had been selectively bred for 14 generations. Prior to testing, zebrafish were housed in 40 L mixed-sex tanks on a custom-built recirculating water system with solid and biological filtration. System water was held at approximately 27°C with a 700 μS conductivity and a 7.2 pH. Zebrafish were on a 14:10 light/dark cycle and fed twice daily with Tetramin Tropical Flakes (Tetra, USA). All procedures and experiments were approved by the Institutional Animal Care and Use Committee of the University of Nebraska at Omaha (17-070-09-FC).

### Caffeine Treatment

We treated proactive and reactive zebrafish with caffeine or vehicle control for 15 minutes. Sample size included 30 proactive zebrafish (n=15 control with 6 females and 9 males; n=15 treated with 5 females and 10 males) and 30 reactive zebrafish (n=15 control with 6 females and 9 males; n=15 treated with 7 females and 8 males). For acute treatment, zebrafish were randomly placed in system water with 0 or 50 mg/L caffeine (Sigma Aldrich) for 15 minutes following previously established protocols^16,24,25^. We used 50 mg/L caffeine concentration after conducting a pilot dose-response study where we found 50 mg/L significantly changed stress-related behavior compared to controls in at least one of the strains (See Supplementary material).

### Behavioral Assay

After 15 minutes of caffeine or vehicle control treatment, we subjected individual zebrafish to a novelty stressor with a novel tank diving test, following established procedures^25,27,28,30,51^. In brief, we used a clear 3-L trapezoidal tank (15.2 height x 28.3 top x 22.5 bottom x 11.4 cm width; Pentair Aquatic Ecosystems) filled with 2-L of system water. We video-recorded the fish for 30 minutes, and the behavior was then quantified using video tracking software (Ethovision Noldus XT Version 17, Wageningen, Netherlands), as previously described^27,29,30,47^. We quantified behavioral indicators of stress for the first 6 minutes, including duration (s) in bottom half of the tank, number of transitions into the top half of the tank, total distance swam (cm), average velocity (cm/s), and time spent frozen (s). The subject was considered frozen if it was moving less than 0.55 cm/s. This time was chosen to ensure we were measuring the response to a novelty stressor without the confounds of habituation^52,53^ and to maintain comparability with similarly used timeframes in many other published studies^16,25,27,28,31^. Increased time frozen and time spent in the bottom half of the tank and decreased transitions to the top half are indicators of heightened stress and anxiety^25,27^. Distance swam and average velocity were used to assess locomotor activity. Behavioral testing occurred between 6-10 hours after light onset. Immediately after the behavior test, we extracted the brains and stored them at -80°C until processing. We determined the sex of each fish by identification of testes or ovaries on dissection.

### Gene Expression Analysis

Using quantitative reverse transcription PCR (qRT-PCR), we quantified the whole-brain gene expression of adenosine receptors (*adora1, adora2aa, adora2ab, adora2b*) and enzymes adenosine deaminase (*ada*) and ecto-5’-nucleotidase (*nt5e*) with *ef1a* used as an endogenous reference for normalization following established protocols^28,29,32^. Expression of *ef1a* is stable across sex, developmental stage, chemical treatment, and tissue type in zebrafish^54^. In brief, we homogenized the brain tissue using zirconium oxide beads (Bullet Blender, Next Advanced) and then extracted and purified RNA (RNeasy Plus Mini, Qiagen). For each sample, we quantified RNA concentration with a Qubit 3.0 (Life Technologies). For each fish, we used 230 ng of total RNA for cDNA synthesis (SuperScript IV First-Strand Synthesis System, Invitrogen) and purified the cDNA using Amicon Ultracentrifugal filters (Millipore).

For qRT-PCR, we utilized the QuantStudio 7 Flex Real-Time PCR System (Applied Biosystems) using SYBR green detection chemistry (PowerUp SYBR Green Master Mix, Applied Biosystems). We designed all primers on Primer-Blast (NCBI), except for *ef1a*, which was determined by past studies^55,56^. The primer concentration was 5 pmol for every gene except *adora2aa* (1 pmol) and *adora2ab* (2.5 pmol) (Supplemental Table S1). Reaction parameters for each gene were as follows: 2 min at 50°C, 2 minutes at 95°C, and 40 cycles of 95°C for 15 sec and 60°C for 1 minute. We ran each sample in triplicate and quantified expression using the relative standard curve method.

### Statistical Analysis

Of the twelve dependent variables, four were not normally distributed even after a log transformation. Therefore, we applied a generalized linear model (GLZM) with the identity link function in SPSS (Version 29) to examine changes in behaviors and normalized gene expressions. Using the GLZM, we investigated main effects of strain (reactive, proactive), sex (female, male), treatment (50 mg/L caffeine, control), strain*treatment interaction, and strain*treatment*sex interaction on the behavior and gene expression. To evaluate the direction of effects, we examined the simple main effects within each GLZM and applied a Benjamini-Hochberg correction when examining the interaction effects^57^. Nonparametric (Spearman’s) correlations were also analyzed between each dependent variable, followed by a Benjamini-Hochberg correction^57^. In addition, we used principal component analysis (PCA) with the five behavioral variables to calculate a composite stress behavior score (principal component scores) for each individual. This score was also used as a dependent variable in the GLZM as described above.

## Results

### Significant effect of strain and sex on composite behavioral scores

Principal component analysis of the five behavioral variables revealed that all behaviors loaded onto a single component (stress behavior axis) where swim velocity, distance swam, and number of transitions into the top half of the tank loaded positively while time frozen and time in the bottom half of the tank loaded negatively. Resulting principal component (PC) scores demonstrated that more negative scores indicate higher stress-related behaviors. There was a significant main effect of strain on the PC scores (Wald *χ*^2^ = 40.389, p = 2.081×10^-10^) with reactive fish having significantly lower scores than proactive fish (Figure 1). There was also a significant main effect of sex on the PC scores (Wald *χ*^2^ = 13.997, p =.000183) with female fish having significantly lower PC scores than males (Figure 1). There was a trend of a main effect of treatment on the PC scores (Wald *χ*^2^ = 3.774, p =.052), in which caffeine treated fish had an increase in PC scores relative to controls. There was also a trend of a strain by treatment by sex interaction effect on the PC scores (Wald *χ*^2^ = 7.660, p =.054). Caffeine treated reactive females had significantly lower PC scores than caffeine treated reactive males (p = 2.84×10^-5^) and caffeine treated proactive females (p = 2.21×10^-4^). Caffeine treated reactive males had significantly higher PC scores than male reactive controls (p =.00368). Female reactive controls had significantly lower PC scores than female proactive controls (p = 7.38×10^-5^). Male reactive controls had significantly lower PC scores than male proactive controls (p = 6.58×10^-4^). There was not a significant strain by treatment interaction effect (Wald *χ*^2^ = 0.986, p =.321).

**Figure 1.**
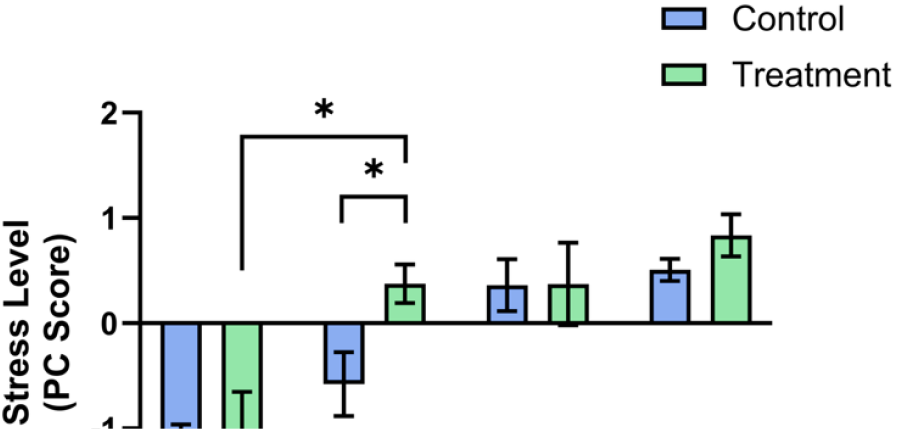
Acute caffeine treatment significantly altered behavior between strains and sexes. For each group, we calculated a stress PC score from distance swam, average velocity, time spent frozen, number of top transitions, and time spent in bottom half. Data shown are mean ± 1 SEM. Significant strain and treatment differences are indicated by an asterisk (*p* ≤ 0.05).

### Stress coping style, acute caffeine exposure, and sex alter discrete behaviors

There were significant main effects of strain on the number of transitions into the top half (Wald *χ*^2^ = 37.667, p = 8.39×10^-10^), time spent frozen (Wald *χ*^2^ = 42.802, p = 6.06×10^-11^), swimming velocity (Wald *χ*^2^ = 28.395, p = 9.89×10^-8^), and distance swam (Wald *χ*^2^ = 28.768, p = 8.16×10^-8^). Compared to proactive fish, reactive fish had less transitions to the top of the tank, spent more time frozen, swam with lower velocity, and swam less total distance (Figure 2). There was not a significant main effect of strain on time spent in the bottom half (Wald *χ*^2^ = 0.107, p =.743).

**Figure 2.**
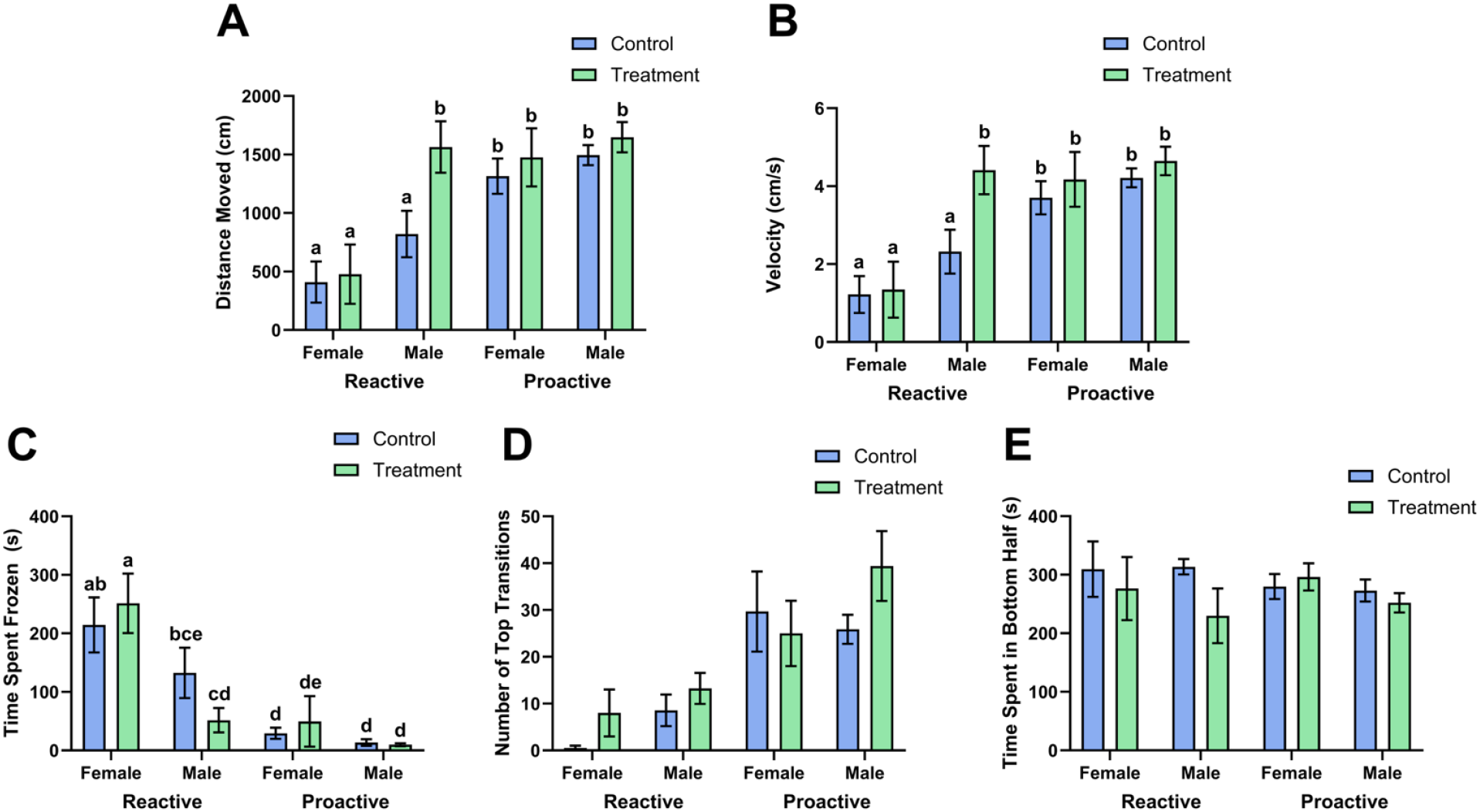
Female zebrafish exhibited higher anxiety-related behavior than males. We measured distance swam (A), average velocity (B), time spent frozen (C), number of top transitions (D), and time spent in bottom half (E) for each group. Data shown are mean ± 1 SEM. Significant differences between groups are indicated by different lower-case letters (*p* ≤ 0.05 after BH corrections).

Caffeine treatment significantly increased swimming velocity (Wald *χ*^2^ = 5.007, p =.025) and distance swam (Wald *χ*^2^ = 5.121, p =.024) compared to control fish. There were no significant main effects of caffeine treatment on time spent in the bottom half (Wald *χ*^2^ = 1.985, p =.159), number of transitions into the top half (Wald *χ*^2^ = 2.074, p =.150), or time spent frozen (Wald *χ*^2^ = 0.107, p =.744). There were no significant strain by treatment interaction effects on behavioral variables [number of transitions into the top half (Wald *χ*^2^ = 0.053, p =.819), time spent in the bottom half (Wald *χ*^2^ = 1.704, p =.192), time spent frozen (Wald *χ*^2^ = 0.523, p =.469), swimming velocity (Wald *χ*^2^ = 0.884, p =.347), and distance swam (Wald *χ*^2^ = 1.010, p =.315)].

There were significant main effects of sex on time spent frozen (Wald *χ*^2^ = 16.247, p = 5.556×10^-5^), swimming velocity (Wald *χ*^2^ = 13.604, p = 2.257×10^-4^), and distance swam (Wald *χ*^2^ = 13.908, p = 1.920×10^-4^) but not on the time spent in the bottom half (Wald *χ*^2^ = 1.178, p =.278) and the number of transitions into the top half (Wald *χ*^2^ = 2.683, p =.101). Compared to male fish, female fish spent more time frozen, swam with less velocity, and swam less distance total (Figure 2).

There were significant strain by treatment by sex interaction effects for time spent frozen (Wald *χ*^2^ = 11.738, p =.008), swimming velocity (Wald *χ*^2^ = 9.386, p =.025), and distance swam (Wald *χ*^2^ = 9.251, p =.026). Caffeine treated reactive females spent significantly more time frozen, swam less distance, and swam at a lower velocity than caffeine treated reactive males (p = 1.02×10^-6^, p = 7.32×10^-6^, p = 6.93×10^-6^, respectively) and caffeine treated proactive females (p = 1.29×10^-5^, p = 2.74×10^-4^, p = 2.47×10^-4^, respectively). Female reactive controls spent significantly more time frozen, lower swimming velocity, and swam a lower distance than female proactive controls (p = 4.93×10^-5^, p =.00110, p = 8.10×10^-4^, respectively). Female reactive controls spent significantly more time frozen than male reactive controls (p =.0491) but did not remain significant after a Benjamini-Hochberg adjustment. Male reactive controls had significantly more time frozen, lower swimming velocity, and swam a lower distance than male proactive controls (p =.00139, p =.00229, p =.00224, respectively). In addition, male reactive controls had significantly lower swimming velocity and swam a lower distance than caffeine treated reactive males (p =.00108, p =.00109, respectively). Male reactive controls had significantly higher time frozen than caffeine treated reactive males (p = 0.0350) but did not remain significant after a Benjamini-Hochberg adjustment. There was not a significant strain by treatment by sex interaction effect on time spent in the bottom half (Wald *χ*^2^ = 1.079, p =.782) and number of transitions into the top half (Wald *χ*^2^ = 3.139, p =.371).

### Differential gene expression of adora1b, adora2aa, adora2ab, adora2b, and ada between zebrafish strains

There were significant main effects of strain on expression of *adora1b* (Wald *χ*^2^ = 13.079, p =.0003), *adora2aa* (Wald *χ*^2^ = 11.075, p =.00088), *adora2ab* (Wald *χ*^2^ = 6.862, p =.0088), *adora2b* (Wald *χ*^2^ = 5.521, p =.019), and *ada* (Wald *χ*^2^ = 4.887, p =.027) but not *nt5e* (Wald *χ*^2^ = 1.983, p =.159). Compared to proactive fish, reactive fish had lower expression of *adora2aa* and higher expression of *adora1b, adora2ab, adora2b*, and *ada* (Figure 3).

**Figure 3.**
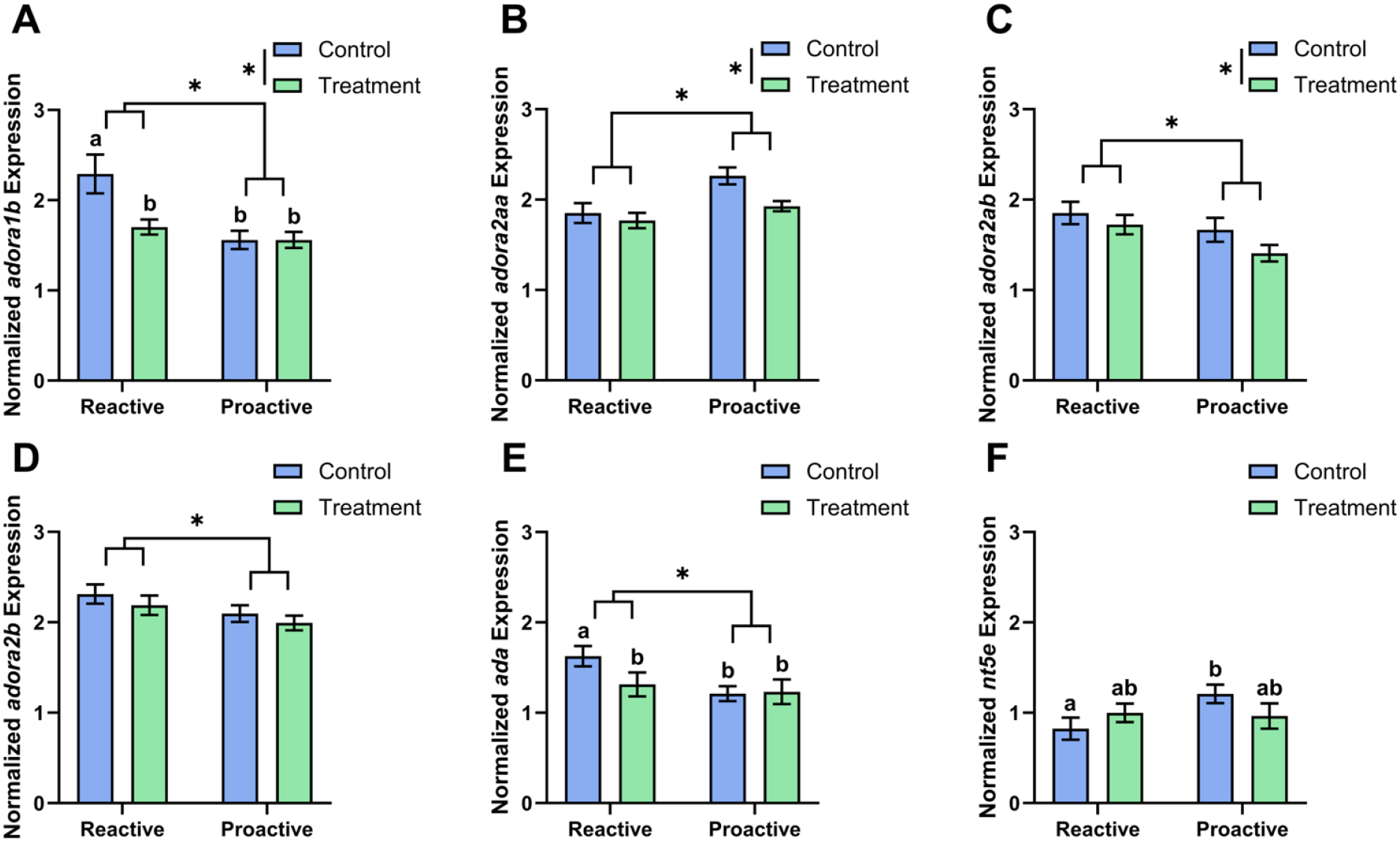
Differential gene expression of *adora1b, adora2aa, adora2ab, adora2b*, and *ada* between zebrafish strains. We measured expression of *adora1b* (A), *adora2aa* (B), *adora2ab* (C), *adora2b* (D), *ada* (E), and *nt5e* (F) for each group. Data shown are mean ± 1 SEM. Significant strain and treatment differences are indicated by an asterisk (*p* ≤ 0.05), and differences between groups are indicated by different lower-case letters (*p* ≤ 0.05 after BH corrections).

There were significant main effects of caffeine treatment on expression of *adora1b* (Wald *χ*^2^ = 5.015, p =.025), *adora2aa* (Wald *χ*^2^ = 4.976, p =.026), and *adora2ab* (Wald *χ*^2^ = 4.381, p =.036) but not adora2b (Wald *χ*^2^ = 1.078, p =.299), *ada* (Wald *χ*^2^ = 1.469, p =.226), or *nt5e* (Wald *χ*^2^ = 0.084, p =.771). Caffeine treated fish had lower expression of *adora1b, adora2aa*, and *adora2ab* compared to control fish (Figure 3).

There was a significant strain by treatment interaction effect for *adora1b* expression (Wald *χ*^2^ = 4.952, p =.026). Reactive control fish had significantly higher expression of *adora1b* than proactive control fish (p = 3.27×10^-5^) and reactive treated fish (p =.00136). There was a trend in strain by treatment interaction effects for expression of *ada* (Wald *χ*^2^ = 3.385, p =.066) and *nt5e* (Wald *χ*^2^ = 2.794, p =.095). Reactive controls had significantly higher expression of *ada* than proactive controls (p =.00398). Reactive controls had significantly higher expression of *ada* than caffeine treated reactive fish (p =.0286) but this did not remain significant after a Benjamini-Hochberg adjustment. Reactive controls had significantly lower expression of *nt5e* than proactive controls (p =.0285) but this did not remain significant after a Benjamini-Hochberg adjustment. There were no significant strain by treatment interaction effects for expression of *adora2aa* (Wald *χ*^2^ = 0.994, p =.319), *adora2ab* (Wald *χ*^2^ = 0.334, p =.563), and *adora2b* (Wald *χ*^2^ = 0.057, p =.811). There was not a significant main effect of sex on gene expression (*adora1b*: Wald *χ*^2^ = 2.444, p =.118; *adora2aa*: Wald *χ*^2^ = 0.159, p =.691; *adora2ab*: Wald *χ*^2^ = 0.001, p =.977; *adora2b*: Wald *χ*^2^ = 0.583, p =.445; *ada*: Wald *χ*^2^ = 1.324, p =.250; *nt5e*: Wald *χ*^2^ = 0.172, p = 0.678).

There was a significant strain by treatment by sex interaction effect for *adora2ab* expression (Wald *χ*^2^ = 10.152, p =.017). Female reactive control fish had significantly higher expression of *adora2ab* than male reactive control fish (p =.0104). Female reactive control fish had significantly higher expression of *adora2ab* than female reactive treated fish (p =.0053) and female proactive controls (p =.023) but this did not remain significant after a Benjamini-Hochberg adjustment. Caffeine treated reactive males had significantly higher expression of *adora2ab* than caffeine treated proactive males (p =.009) but this did not remain significant after a Benjamini-Hochberg adjustment.

There was a trend in the strain by treatment by sex interaction effect for expression of *adora2aa* (Wald *χ*^2^ = 7.176, p =.067). Male reactive controls had significantly lower expression of *adora2aa* than male proactive controls (p = 1.99×10^-5^). Male proactive controls had significantly higher expression of *adora2aa* than caffeine treated proactive males (p = 1.66×10^-4^). Female proactive controls had a trend of higher expression of *adora2aa* than male proactive controls (p =.0448) but did not remain significant after a Benjamini-Hochberg adjustment. There were no significant strain by treatment by sex interaction effects for expression of *adora1b* (Wald *χ*^2^ = 0.811, p =.847), *adora2b* (Wald *χ*^2^ = 4.132, p =.248), *ada* (Wald *χ*^2^ = 5.481, p =.140), and *nt5e* (Wald *χ*^2^ = 1.007, p =.799).

### Expression of adora2ab and ada are positively correlated with anxiety behavior

There were significant correlations between the PC score and the expression of *adora2ab* (*p* =.003, ρ = -.375) and *ada* (*p* =.001, ρ = -.410), in which the higher the expression, the higher the anxiety behavior (Figure 4). While there was a significant correlation between *adora2b* expression and PC score (*p* =.028, ρ = -.285), this relationship did not remain significant after a Benjamini-Hochberg adjustment. There was not a significant correlation between the PC score and expression of *adora1b* (*p* =.0916, ρ = -.220), *adora2aa* (*p* =.174, ρ = 0.178), or *nt5e* (*p* =.503, ρ =.088).

**Figure 4.**
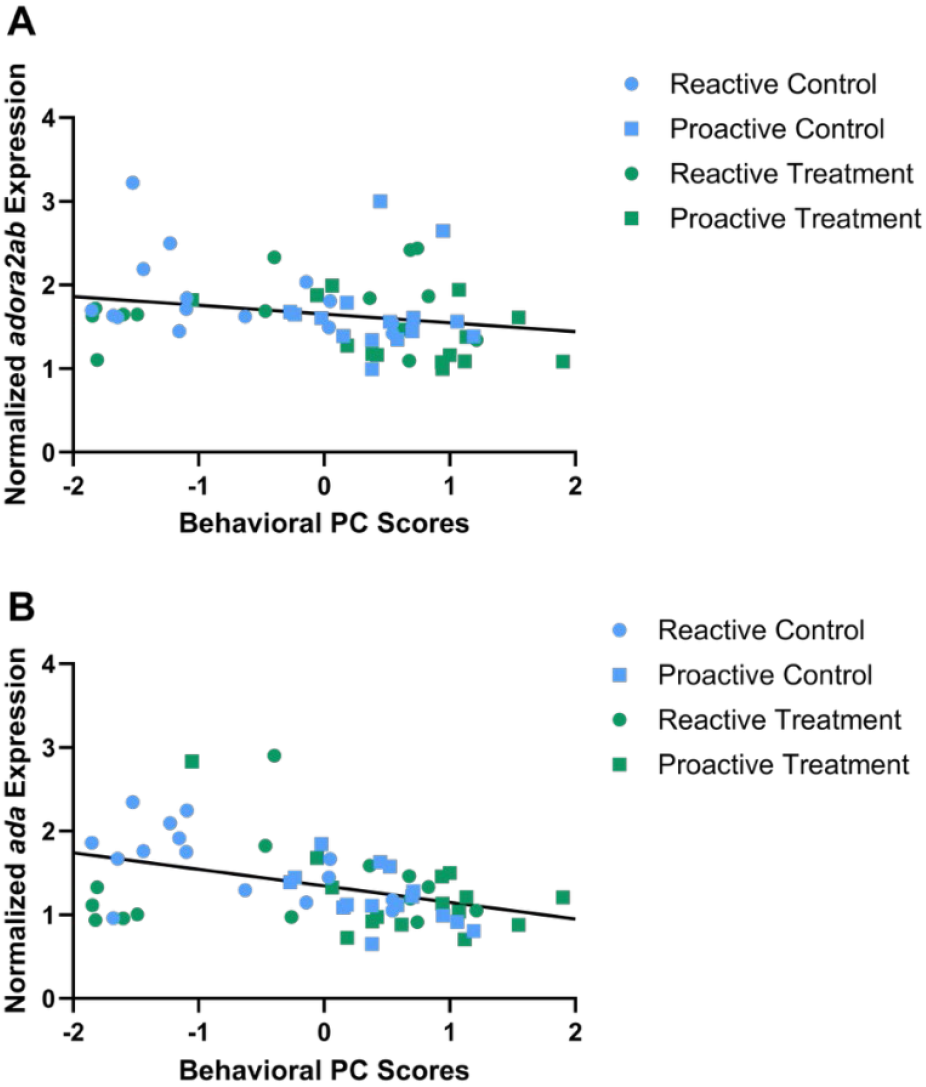
Expression of *adora2ab* and *ada* are correlated with the composite stress behavior scores. Significant correlation between gene expression, *adora2ab* (A) and *ada* (B), and behavioral PC scores within all zebrafish (reactive control, blue circle; proactive control, blue square; reactive treatment, green circle; proactive treatment, green square).

## Discussion

This study investigated how the adenosine signaling pathway and antagonism of the pathway with acute caffeine treatment influences behavior of proactive and reactive zebrafish strains. We found that reactive zebrafish express higher levels of adenosine receptors A1B, A2Ab, A2B, and adenosine deaminase and lower levels of adenosine receptor A2Aa than proactive zebrafish. In addition, acute caffeine exposure differentially affected the stress coping styles and the sexes. The proactive strain behavior was not significantly impacted by caffeine treatment while the reactive strain behavior differed by sex. Following caffeine treatment, female reactive zebrafish exhibited higher anxiety behavior and male reactive zebrafish exhibiting lower anxiety but hyperactive behavior. We also noted that overall female zebrafish had higher anxiety behavior and demonstrated that *adora2ab* and *ada* expression is positively correlated with anxiety behavior. These findings indicate that variation in adenosine signaling may contribute to differences in anxiety-related behavior between the stress coping styles and sexes.

Adenosine A1 receptors are inhibitory and widespread through the brain while adenosine A_2A_ receptors are excitatory and have the highest density in the basal ganglia, amygdala, and hypothalamus. Adenosine receptor expression levels within the brain are associated with the anxiety response. Low anxiety rats were found to have high A1R and low A2AR expression in the ventral hippocampus, which favors the inhibitory effect of A1R. High anxiety rats expressed low levels of A1R, favoring the excitatory effect of A2AR, increasing glutamate release and causing activation of the hypothalamic-pituitary-adrenal (HPA) axis^58^. Our research supports that an imbalance between the inhibitory A1R and the excitatory A2AR expression may impact anxiety-like behavior. Previous research supports that the adenosine A2A receptor is closely linked with anxiety-related behavior in humans and animal models^12–17,20–23,58^. Our research in zebrafish does not show the same pattern between A1AR and A2AR but suggests that the A2Aa and A2Ab receptors have counteracting effects with A2Ab being more highly expressed in the high stress reactive strain and A2Aa being more highly expressed in the low stress proactive strain. We also found that A2Ab expression, but not A2Aa expression, is highly correlated with anxiety behavior across all tested zebrafish.

Behavioral effects of adenosine receptor expression levels have not been fully investigated. Overexpression of A2A receptors in rats causes depressive-like behavior and hyperlocomotion^59^. Gene expression of A1b, A2Aa, A2Ab, and A2B increased in the zebrafish brain after an acute restraint stressor^60^. This study suggests that A2Aa and A2Ab expression increased due to stress, and A1b expression increased as a compensatory mechanism to maintain equilibrium of excitatory signals between the A2A and A1 receptors. The contrasting A2Aa and A2Ab receptor expression in the zebrafish brain has not been fully investigated but has been noted in previous studies. Following exposure to methionine, mRNA levels of A1 and A2Aa increased while A2Ab levels decreased^61^. Expression of A2Aa increased after exposure to valproic acid while A2Ab expression did not change^62^. The results from these studies suggest that A2Aa and A2Ab receptor expression may be controlled through separate mechanisms and respond to stressors in distinct ways.

In this study, caffeine treatment decreased expression of *adora1b, adora2aa*, and *adora2ab*, independent of strain. This is consistent with the known pharmacodynamics of caffeine as an adenosine receptor antagonist, which can lead to downregulation of receptor mRNA expression through feedback mechanisms^63–65^. The suppression of these genes supports the interpretation that caffeine engaged its molecular targets, leading to behavioral differences. Previous studies have shown that double heterozygous A1R and A2AR mice portray similar effects of chronic caffeine use (5-7 days)^66^, suggesting that the downregulation of A1 and A2A receptors may be partially responsible for the effects of caffeine. It is also of note that the adenosine receptor downregulation in the current study occurred relatively quickly, within an hour of acute caffeine treatment.

Adenosine levels are closely regulated through complex processes, including metabolism and transport. Expression and activity of enzymes within the adenosine signaling pathway can impact adenosine concentrations and alter behavioral states. Adenosine deaminase, which degrades adenosine into inosine, has been linked to anxiety. Adenosine deaminase knockout mice have increased anxiety behavior^67^. Unpredictable chronic stress has been found to decrease adenosine deaminase activity and increase extracellular adenosine levels in zebrafish brains, potentially acting as a calming mechanism^10^. Our study found that adenosine deaminase expression is correlated with anxiety behavior, in which high expression is associated with high anxiety. A potential mechanism linking strain differences in adenosine signaling and behavior may be that the reactive strain has reduced adenosine levels due to elevated expression of adenosine deaminase, which degrades adenosine, and slightly reduced expression of ecto-5’-nucleotidase, which produces it. The high anxiety behavior observed within the reactive zebrafish strain may partially be caused by low adenosine levels. In addition, caffeine treatment likely acts as a stressor, indicated by increased whole-body cortisol in zebrafish following exposure^16,24,68^. We have shown that the caffeine-treated reactive zebrafish had significantly lower adenosine deaminase expression than controls, potentially indicating an increase in adenosine that may act as a calming mechanism in response to caffeine exposure.

We found significant main effects of strain, treatment, and sex on a composite measure of stress behavior. Consistent with prior studies, the reactive strain and females had increased stress behaviors^27–31^. Surprisingly, we observed that caffeine treatment resulted in a reduction of stress behaviors. While many prior studies have documented anxiogenic effects of caffeine, this relationship is complex^14–17,24,25,69^. Strain- and caffeine concentration-specific behavioral effects in response to acute caffeine exposure have been noted in prior studies^16,17^. When investigating the behavioral response to caffeine in wild type and leopard zebrafish populations, the leopard zebrafish had higher baseline cortisol levels and exhibited heighted anxiety responses to caffeine compared to wild type zebrafish^16^. In addition, low dose caffeine typically induces stimulatory effects, indicated by increased swimming distance and speed^13,25,50,70^. On the other hand, high doses of caffeine commonly induce anxiety behavior, indicated by decreased swimming distance and increased time frozen^13,25,50^.

Intriguingly, we found a significant strain by treatment by sex interaction effect on the composite measure of stress behavior. The anxiolytic effect of caffeine treatment was only seen in males with the reactive stress coping style. This strain and sex specific effect of caffeine was also seen when examining discrete behaviors where female reactive fish became more anxious after caffeine treatment, while reactive male fish became less anxious but more hyperactive. Our results emphasize the importance of accounting for stress coping style and sex when evaluating the impact of adenosine signaling modulators. Further, it suggests that biases in behavioral stress response characteristic of each stress coping style may be primarily driven by adenosine neuromodulation in males with the reactive stress coping style. Other studies have found strain-specific effects of a drug (including ethanol, caffeine, cannabidiol)^16,17,31,71^. In addition, sex-specific behavior in response to caffeine has been seen previously in zebrafish^24^, in which acute 300 mg/L caffeine exposure caused female fish to have more freezing behavior and higher whole-body cortisol compared to male fish, while male fish had increased erratic movement and higher testosterone compared to female fish. In addition, gene ontology pathway analysis shows that caffeine may impact steroid hormone biosynthesis, indicating that caffeine may differentially affect stress behavior between the sexes due to altered cortisol and testosterone production^24^. On the other hand, some studies did not find differential effects of sex on anxiety behavior following caffeine treatment^17,72^. To our knowledge, this is the first study showing both strain- and sex-dependent effects of caffeine.

## Conclusion

Overall, our findings reinforce the central role of coping style in shaping both behavioral and molecular responses to stress and neuromodulatory compounds. Acute caffeine exposure exerts coping style- and sex-dependent effects on anxiety-like behavior in zebrafish, with reactive individuals, particularly males, showing the greatest sensitivity. The accompanying changes in adenosine receptor and adenosine deaminase expression demonstrate a possible molecular mechanism underlying these differences, highlighting the adenosine signaling pathway as a promising target for understanding individual variability in anxiety. We hypothesize that a combined effect of adenosine receptor expression, extracellular adenosine levels, and downstream adenosine signaling contributes to how the adenosine signaling pathway impacts anxiety-related behavior. Further, this mechanism can explain, in part, the different biases in behavioral and physiological responses to stress between the stress coping styles. Future studies incorporating region-specific brain analyses, chronic caffeine exposure paradigms, or more targeted pharmacological manipulations will help clarify the causal links between adenosine signaling, stress coping style, and anxiety behavior.

## Acknowledgements

We are grateful to the UNO Animal Care and Use Program and members of the Wong laboratory for help with zebrafish husbandry. We thank Levi Storks, Jamie Corcoran, Abigail Reynolds, and other members of the Wong lab for helpful discussions and technical assistance. This study was supported by the UNO Fund for Undergraduate Scholarly Experiences to S.E.K. and the National Institutes of Health (R15MH113074) to R.Y.W.

## Author Contributions

S.E.K. and R.Y.W. conceived the study, conducted statistical analyses, and wrote the manuscript. S.E.K. conducted behavioral testing, brain extraction, gene expression quantification, and data collection.

## Data Availability Statement

All data generated or analyzed during this study are included in this published article and its Supplementary Information files.

