## Supplemental Information for "Differential adenosine signaling and effects of acute caffeine exposure on alternative stress coping styles in zebrafish (*Danio rerio*)"

Author Affiliation: Department of Biology, University of Nebraska at Omaha

**Supplemental Methods and Results**

To identify a biologically relevant dose and behavioral trial length applicable to both proactive and reactive zebrafish, we acutely treated with 0 (n = 16 of each coping style), 50 (n = 8 of each coping style), 100 (n = 8 of each coping style), or 150 mg/L caffeine concentrations (n = 8 of each coping style) for 15 minutes and then quantified stress behaviors (distance swam, time spent in bottom half of tank, swimming velocity, time frozen) in the novel tank diving test for 6 minutes. We used a generalized linear model (GLZM) in SPSS (Version 29) to examine main effects of strain (reactive, proactive), caffeine concentration (0, 50, 100, 150 mg/L caffeine) and strain by treatment interaction effect on distance swam, time spent in bottom half of tank, swimming velocity. For time frozen we used GLZM to examine the main effects of strain (reactive, proactive), caffeine concentration (0, 50 mg/L caffeine) and strain by treatment interaction effect. To evaluate the direction of effects, we examined the simple main effects within each GLZM. Overall, caffeine concentration had minimal effects on stress behavior of proactive zebrafish. However, in the reactive strain, exposure to 50 mg/L caffeine increased stress behavior and 100 mg/L caffeine increased locomotion relative to controls.

There was no main effect of caffeine concentration on distance swam (Wald $\chi^2$ = 5.765, p = .124), time spent in the bottom half (Wald $\chi^2$ = 3.863, p = .277), swimming velocity (Wald $\chi^2$ = 5.682, p = .128), and time spent frozen (Wald $\chi^2$ = 2.122, p = .145). There were significant main effects of strain on distance swam (Wald $\chi^2$ = 114.515, p < .001), time spent in the bottom half (Wald $\chi^2$ = 30.857, p = 2.7 * $10^{-8}$), swimming velocity (Wald $\chi^2$ = 114.453, p < .001), and time spent frozen (Wald $\chi^2$ = 118.24, p < .001). There were significant strain by treatment interaction effects on distance swam (Wald $\chi^2$ = 9.045, p = .029), swimming velocity (Wald $\chi^2$ = 8.706, p = .033), and time spent frozen (Wald $\chi^2$ = 5.038, p = .025). Reactive fish treated with 50 mg/L caffeine spent significantly more time frozen than reactive controls (p = .009). There was a trend for reactive fish treated with 50 mg/L caffeine to swim less distance (p = .122) and with slower velocity (p = .138) than reactive controls. There were no significant differences between 50 mg/L caffeine treated and control proactive fish for distance swam (p = .277) and swim velocity (p = .282). Reactive fish treated with 100 mg/L caffeine swam significantly higher distance (p = .014) and with faster swimming velocity (p = .015) than reactive controls. Treatment of 150 mg/L caffeine did not significantly impact the behavior of reactive fish. Neither 100 mg/L nor 150 mg/L caffeine treatment significantly impacted the behavior of proactive fish. There was not a significant strain by treatment interaction effect on time spent in the bottom half (Wald $\chi^2$ = 5.525, p = .137).

| Gene | Primer Conc. | Amplicon Length (bp) | Forward Primer | Reverse Primer |
|---|---|---|---|---|
| *adora1b* | 5 pmol | 95 | 5’-GTCCAGTCATTCGGAACCCA-3’ | 5’-GCCAGCTAATTGCGAACAGG-3’ |
| *adora2aa* | 1 pmol | 121 | 5’-GTTGCCTGTTCATCGCCTG-3’ | 5’-ACTAGGCTGTTGTACCTAAGCG-3’ |
| *adora2ab* | 2.5 pmol | 89 | 5’-CTTTTCGCAGTTTGCTGGCT-3’ | 5’-ATGACCCAATCCTGAGGTCG-3’ |
| *adora2b* | 5 pmol | 76 | 5’-AGTCTTCTAGCGGTTGCCAT-3’ | 5’-TACCTGTGACCAGCTCCCTA-3’ |
| *ada* | 5 pmol | 134 | 5’-CGGCGAGGAATCAGTCTACC-3’ | 5’-TCTCTGTCCCCTGCAATGAC-3’ |
| *nt5e* | 5 pmol | 117 | 5’-CTGGTCCCCAACAAGGTGTA-3’ | 5’-TATATCCAGATCACCGCTGTCG-3’ |
| *ef1a* | 5 pmol | 150 | 5’-CCTCTTGGTCGCTTTGC-3’ | 5’-GGTGTGATTGAGGGAAATTCA-3’ |

**Supplemental Table S1. qRT-PCR Primers**

| ID | Strain | Treatment | Sex | PC Score | Distance Swam (cm) | Velocity (cm/s) | Time Frozen (s) | Top Transitions | Bottom Time (s) | ***adora1b / ef1a*** | ***adora2aa / ef1a*** | ***adora2ab / ef1a*** | ***adora2b/ ef1a*** | ***ada/ ef1a*** | ***nt5e/ ef1a*** |
|---|---|---|---|---|---|---|---|---|---|---|---|---|---|---|---|
| R1C | Reactive | Control | F | -1.525 | 163.89 | 0.46 | 240.94 | 0 | 360.00 | 2.23 | 2.20 | 2.60 | 2.60 | 2.35 | 1.34 |
| R2C | Reactive | Control | F | -1.440 | 34.99 | 0.49 | 336.71 | 0 | 73.91 | 2.10 | 2.31 | 2.19 | 1.99 | 1.76 | 0.54 |
| R3C | Reactive | Control | M | -1.852 | 70.30 | 0.20 | 353.39 | 0 | 360.00 | 4.60 | 1.35 | 1.70 | 2.00 | 1.86 | 1.60 |
| R4C | Reactive | Control | M | -1.094 | 446.81 | 1.26 | 151.99 | 0 | 360.00 | 1.58 | 1.10 | 1.84 | 1.32 | 2.25 | 0.39 |
| R5C | Reactive | Control | M | 0.543 | 1532.19 | 4.31 | 10.21 | 28 | 294.74 | 3.10 | 1.89 | 1.54 | 2.61 | 1.05 | 0.02 |
| R6C | Reactive | Control | F | -1.680 | 165.48 | 0.47 | 311.42 | 0 | 360.00 | 2.62 | 1.36 | 1.63 | 1.94 | 0.96 | 0.62 |
| R7C | Reactive | Control | M | -1.647 | 134.56 | 0.38 | 284.73 | 0 | 360.00 | 1.89 | 1.57 | 1.61 | 2.52 | 1.67 | 1.26 |
| R8C | Reactive | Control | M | 0.046 | 1218.61 | 3.44 | 12.08 | 12 | 325.60 | 2.55 | 1.63 | 1.81 | 2.54 | 1.67 | 0.59 |
| R9C | Reactive | Control | F | -0.143 | 1225.91 | 3.45 | 13.95 | 0 | 360.00 | 2.73 | 2.17 | 2.04 | 3.02 | 1.15 | 0.81 |
| R10C | Reactive | Control | F | -1.227 | 403.85 | 1.14 | 196.06 | 0 | 360.00 | 1.70 | 2.12 | 2.50 | 2.23 | 2.10 | 0.96 |
| R11C | Reactive | Control | F | -1.099 | 465.81 | 1.31 | 187.12 | 3 | 343.24 | 1.83 | 1.71 | 1.71 | 2.34 | 1.75 | 1.07 |
| R12C | Reactive | Control | M | 0.546 | 1589.02 | 4.48 | 4.64 | 21 | 270.74 | 1.65 | 1.85 | 1.42 | 2.17 | 1.18 | 0.66 |
| R13C | Reactive | Control | M | -0.629 | 797.24 | 2.25 | 137.54 | 6 | 303.67 | 1.76 | 1.95 | 1.62 | 2.73 | 1.30 | 1.09 |
| R14C | Reactive | Control | M | -1.155 | 340.57 | 0.96 | 203.14 | 2 | 264.77 | 1.23 | 2.75 | 1.45 | 2.52 | 1.92 | 0.05 |
| R15C | Reactive | Control | M | 0.039 | 1258.65 | 3.61 | 34.70 | 8 | 283.45 | 2.80 | 1.81 | 1.49 | 2.13 | 1.45 | 1.35 |
| P1C | Proactive | Control | M | -0.023 | 1020.30 | 2.87 | 15.75 | 18 | 300.91 | 1.93 | 2.08 | 1.60 | 2.26 | 1.84 | 1.53 |
| P2C | Proactive | Control | M | 0.522 | 1509.32 | 4.25 | 5.24 | 24 | 264.87 | 1.56 | 2.74 | 1.56 | 1.95 | 1.57 | 1.37 |
| P3C | Proactive | Control | M | 0.446 | 1254.11 | 3.54 | 10.28 | 27 | 176.02 | 1.90 | 2.93 | 3.00 | 2.92 | 1.63 | 0.90 |
| P4C | Proactive | Control | F | 1.189 | 1846.87 | 5.21 | 5.51 | 55 | 261.57 | 1.33 | 1.88 | 1.39 | 1.94 | 0.80 | 1.34 |
| P5C | Proactive | Control | M | 0.182 | 1349.12 | 3.79 | 57.76 | 22 | 319.49 | 2.13 | 2.06 | 1.79 | 2.24 | 1.12 | 1.65 |
| P6C | Proactive | Control | F | 0.151 | 1386.55 | 3.90 | 46.55 | 17 | 340.01 | 1.49 | 1.93 | 1.39 | 2.04 | 1.09 | 1.43 |
| P7C | Proactive | Control | F | 0.948 | 1439.60 | 4.05 | 4.24 | 57 | 204.78 | 1.87 | 2.35 | 2.65 | 2.34 | 0.99 | 1.65 |
| P8C | Proactive | Control | M | 0.711 | 1778.36 | 5.02 | 5.31 | 25 | 304.18 | 1.94 | 2.37 | 1.61 | 1.91 | 1.28 | 1.71 |
| P9C | Proactive | Control | M | 0.582 | 1434.53 | 4.04 | 10.51 | 35 | 267.14 | 1.72 | 2.44 | 1.35 | 2.29 | 1.11 | 1.60 |
| P10C | Proactive | Control | F | -0.272 | 1047.33 | 2.94 | 61.16 | 8 | 339.08 | 1.40 | 1.85 | 1.68 | 2.32 | 1.39 | 0.66 |
| P11C | Proactive | Control | M | 0.701 | 1704.02 | 4.80 | 5.34 | 30 | 312.39 | 1.59 | 2.24 | 1.44 | 2.17 | 1.22 | 1.00 |
| P12C | Proactive | Control | M | 1.060 | 1771.17 | 4.99 | 4.80 | 42 | 185.03 | 0.93 | 2.12 | 1.56 | 1.76 | 0.92 | 0.67 |
| P13C | Proactive | Control | F | 0.379 | 1406.32 | 3.95 | 20.25 | 23 | 274.52 | 0.71 | 1.70 | 0.99 | 1.23 | 0.65 | 0.97 |
| P14C | Proactive | Control | F | -0.233 | 766.30 | 2.16 | 38.20 | 18 | 259.40 | 1.23 | 2.67 | 1.65 | 2.03 | 1.45 | 0.55 |
| P15C | Proactive | Control | M | 0.381 | 1640.14 | 4.62 | 6.51 | 10 | 326.29 | 1.65 | 2.56 | 1.34 | 2.04 | 1.11 | 1.12 |
| R1T | Reactive | Treated | F | 1.213 | 1901.27 | 5.36 | 3.87 | 36 | 89.26 | 1.38 | 2.47 | 1.34 | 1.79 | 1.05 | 1.33 |
| R2T | Reactive | Treated | F | -1.810 | 87.90 | 0.25 | 340.97 | 0 | 360.00 | 1.57 | 1.60 | 1.10 | 2.79 | 1.33 | 0.73 |
| R3T | Reactive | Treated | F | -1.601 | 143.32 | 0.40 | 267.61 | 0 | 360.00 | 1.79 | 1.63 | 1.65 | 2.02 | 0.96 | 1.06 |
| R4T | Reactive | Treated | F | -1.823 | 77.45 | 0.22 | 342.98 | 0 | 360.00 | 1.38 | 1.01 | 1.72 | 2.18 | 0.94 | 0.02 |
| R5T | Reactive | Treated | M | 0.743 | 1654.62 | 4.66 | 43.38 | 31 | 183.72 | 1.96 | 1.56 | 2.44 | 2.88 | 0.91 | 1.13 |
| R6T | Reactive | Treated | M | 0.686 | 2143.72 | 6.06 | 71.37 | 12 | 321.43 | 1.99 | 1.79 | 2.42 | 2.20 | 1.19 | 0.67 |
| R7T | Reactive | Treated | F | -1.847 | 77.13 | 0.22 | 353.79 | 0 | 360.00 | 2.21 | 2.07 | 1.63 | 2.75 | 1.11 | 0.93 |
| R8T | Reactive | Treated | M | 0.831 | 2204.71 | 6.20 | 1.87 | 10 | 347.75 | 1.89 | 1.96 | 1.86 | 1.59 | 1.33 | 1.41 |
| R9T | Reactive | Treated | M | -0.398 | 839.71 | 2.37 | 134.80 | 11 | 197.66 | 2.31 | 1.92 | 2.33 | 1.98 | 2.90 | 1.39 |
| R10T | Reactive | Treated | M | 0.677 | 2144.21 | 6.07 | 3.47 | 2 | 353.59 | 1.39 | 1.89 | 1.09 | 1.72 | 1.46 | 1.28 |
| R11T | Reactive | Treated | M | 0.361 | 1214.46 | 3.42 | 8.54 | 15 | 98.97 | 1.30 | 1.51 | 1.84 | 1.92 | 1.59 | 1.23 |
| R12T | Reactive | Treated | M | 0.573 | 1722.53 | 4.85 | 6.71 | 21 | 335.07 | 1.62 | 1.52 | 1.43 | 2.12 | 1.13 | 0.53 |
| R13T | Reactive | Treated | F | -1.489 | 313.49 | 0.88 | 320.56 | 7 | 357.83 | 1.49 | 1.74 | 1.65 | 2.58 | 1.00 | 1.00 |

| | | | | | | | | | | | | | | | |
|---|---|---|---|---|---|---|---|---|---|---|---|---|---|---|---|
| R14T | Reactive | Treated | F | -0.260 | 742.26 | 2.09 | 129.60 | 13 | 47.28 | 1.88 | 2.00 | 1.66 | 2.49 | 0.97 | 1.47 |
| R15T | Reactive | Treated | M | -0.469 | 589.71 | 1.67 | 142.58 | 4 | 0.83 | 1.36 | 1.88 | 1.69 | 1.83 | 1.82 | 0.81 |
| P1T | Proactive | Treated | M | 0.381 | 1306.00 | 3.68 | 8.88 | 25 | 246.32 | 1.44 | 1.82 | 1.17 | 2.13 | 0.92 | 0.85 |
| P2T | Proactive | Treated | M | 0.062 | 1392.04 | 3.93 | 14.21 | 5 | 355.33 | 1.44 | 1.87 | 1.99 | 1.80 | 1.32 | 2.14 |
| P3T | Proactive | Treated | F | 1.001 | 1760.51 | 5.05 | 5.64 | 40 | 215.05 | 1.10 | 2.40 | 1.16 | 2.50 | 1.50 | 1.94 |
| P4T | Proactive | Treated | M | 0.942 | 1794.09 | 5.07 | 5.27 | 30 | 176.11 | 1.41 | 1.76 | 1.00 | 1.77 | 1.13 | 0.80 |
| P5T | Proactive | Treated | M | -0.058 | 845.57 | 2.38 | 19.29 | 23 | 254.80 | 2.09 | 1.88 | 1.88 | 2.38 | 1.68 | 1.01 |
| P6T | Proactive | Treated | M | 1.548 | 2070.69 | 5.83 | 4.30 | 63 | 202.24 | 1.56 | 1.85 | 1.61 | 1.99 | 0.88 | 0.73 |
| P7T | Proactive | Treated | M | 1.072 | 1950.08 | 5.49 | 3.50 | 35 | 226.83 | 1.65 | 1.85 | 1.94 | 2.59 | 1.04 | 0.77 |
| P8T | Proactive | Treated | F | 0.616 | 1532.23 | 4.31 | 5.47 | 33 | 294.27 | 1.28 | 1.81 | 1.47 | 1.87 | 0.88 | 0.61 |
| P9T | Proactive | Treated | F | 1.118 | 2146.56 | 6.04 | 4.54 | 33 | 301.24 | 1.63 | 1.71 | 1.09 | 2.11 | 0.71 | 0.54 |
| P10T | Proactive | Treated | M | 0.420 | 1438.01 | 4.07 | 26.73 | 29 | 314.56 | 0.92 | 1.99 | 1.16 | 1.49 | 0.97 | 0.79 |
| P11T | Proactive | Treated | F | 0.183 | 1272.06 | 3.59 | 10.74 | 18 | 312.65 | 1.24 | 2.37 | 1.28 | 1.92 | 0.73 | 0.55 |
| P12T | Proactive | Treated | F | -1.054 | 664.26 | 1.88 | 221.92 | 1 | 358.33 | 1.77 | 2.06 | 1.82 | 1.76 | 2.83 | 1.19 |
| P13T | Proactive | Treated | M | 0.939 | 1823.00 | 5.14 | 4.77 | 37 | 269.14 | 1.99 | 2.04 | 1.08 | 1.96 | 1.46 | 1.29 |
| P14T | Proactive | Treated | M | 1.133 | 1656.22 | 4.67 | 8.01 | 62 | 245.11 | 1.85 | 1.90 | 1.38 | 2.07 | 1.21 | 0.03 |
| P15T | Proactive | Treated | M | 1.904 | 2206.66 | 6.23 | 2.70 | 85 | 230.04 | 2.01 | 1.61 | 1.08 | 1.55 | 1.21 | 1.22 |

**Supplemental Table S2. Raw Behavior and Gene Expression Data.**
